# Diet induced hyperlipidemia confers resistance to standard therapy in pancreatic cancer by selecting for “tumor protective” microbial metabolites and treatment refractory cells

**DOI:** 10.1101/2021.01.12.426380

**Authors:** Kousik Kesh, Roberto Mendez, Beatriz Mateo-Victoriano, Vanessa T Garrido, Brittany Durden, Vineet K Gupta, Alfredo Oliveras Reyes, Jashodeep Datta, Nipun Merchant, Santanu Banerjee, Sulagna Banerjee

**Author notes:** **Address of Correspondence:** Sulagna Banerjee, PhD, Department of Surgery, University of Miami, Biomedical Research Building, Suite 508, 1501, NW 10^th^ Ave, Miami, FL 33156, Or Santanu Banerjee, PhD, Department of Surgery, University of Miami, Biomedical Research Building, Suite 516, 1501, NW 10^th^ Ave, Miami, FL 33156. Contributed equally (Kousik Kesh, Roberto Mendez).

## Abstract

Obesity causes a number of systemic alterations including chronic inflammation and changes in gut microbiome. However, whether these actively contribute to poor survival and therapy resistance in patients with pancreatic cancer remain undetermined. Our current study shows that high fat diet fed pancreatic tumor bearing mice do not respond to standard of care therapy with gemcitabine and paclitaxel when compared to corresponding control diet fed mice. Upon fecal matter transplant from control mice to high fat diet fed mice, the tumors became sensitive to standard of care therapy and showed extensive cell death. Analysis of gut microbiome showed an enrichment of queuosine (Q) producing bacteria in high fat diet fed mice and an enrichment of S-adenosyl methionine (SAM) producing bacteria in control diet fed mice. Further, treatment of high fat diet fed animals with SAM recapitulated the observation with lean to obese fecal matter transplant. Additionally, treatment of pancreatic and colon cancer cell lines in vitro with Q promoted resistance to the paclitaxel and oxaliplatin respectively, while treatment with SAM promoted sensitivity to these drugs. Treatment of pancreatic cancer cells with Q showed upregulation PRDX1, that is involved in oxidative stress protection. Analysis of tumor tissues in high fat diet fed mice showed high PRDX1, low apoptosis and increased proliferation, which were reversed upon treatment with SAM as well as by lean to obese fecal matter transplant. In parallel, high fat diet fed mice showed increase in CD133+ treatment refractory population compared to the control animals. Interestingly, treatment with Q *in vitro* did not enrich for CD133+ population, indicating that Q mediated protection from cell death was independent of enrichment of treatment refractory cells.

These observations indicated that microbial metabolite Q accumulated in high fat diet fed mice protected tumors from chemotherapy induced oxidative stress by upregulating PRDX1. This protection could be reversed by treatment with SAM. We conclude that relative concentration of S-adenosyl methionine and queuosine in fecal samples of pancreatic cancer patients can be indicative of therapy response in this disease.

## Introduction

Over last few decades, prevalence of overweight and obesity have increased worldwide, making it one of the major risk factors for a number of cancers, including pancreatic cancer (Cascetta et al., 2018; Pothuraju et al., 2018; Rawla et al., 2019). Pancreatic cancer cases have been rising steadily in United States, being the 3^rd^ leading cause of cancer related deaths in United States and predicted to become the 2^nd^ most common cancer by 2030. In 2020 alone, the predicted number of pancreatic cancer patients is over 55,000 with more than 90% succumbing to it. Thus, rising numbers in obesity are being correlated with the increased pancreatic cancer cases. While numerous epidemiological studies have established obesity as a risk for pancreatic cancer, the direct role of obesity in onset, progression and prognosis of pancreatic cancer has not been studied. Studies have shown that the risk of pancreatic ductal adenocarcinoma (PDAC) is about 47% greater for patients with high body mass index (BMI), particularly those with increased abdominal adiposity (Genkinger et al., 2011). Studies also showed that individuals who were obese/overweight in early adulthood, had 54% increased risk for PDAC (Genkinger et al., 2011). These studies indicated that increased body weight led to poor survival in pancreatic cancer patients and increased mortality by 2-fold (Calle et al., 2003; McWilliams et al., 2010). A large-scale study in 2009 showed overweight and obesity was associated with lower overall survival in patients with pancreatic cancer (Li et al., 2009). However, despite these efforts, mechanisms that contribute to poor prognosis, dismal survival and resistance to therapy have remained unknown.

Diet induced obesity affects systemic parameters like inflammation, along with altering the gut microbiome significantly, thereby altering the critical balance between the host and microbial metabolites (Cani and Jordan, 2018). Further, in several cancers including pancreatic cancer, gut microbiome determines tumor progression, response to therapy as well as prognosis (Kesh et al., 2020b; Mendez et al., 2020; Mima et al., 2017; Routy et al., 2018; Zheng et al., 2019). In fact, tumor microbiota has also been shown to contribute to gemcitabine resistance in pancreatic cancer (Geller et al., 2017). While the association of microbiome in tumor progression as well as in determining its properties are being evaluated, there is a lacuna in understanding the mechanism by which bacteria may affect these processes.

The role of microbial metabolites in tumor progression is one aspect of microbiome-cancer research field that has remained underexplored. While some studies show the role of short chain fatty acid (SCFA) in colon cancer progression, their role in cancers that are more remote and not exposed to the microbial milieu is not clear. Certain microbial metabolites like polyamines supplement the host metabolic pool and contribute to tumor cell proliferation by feeding into nucleotide and protein synthesis pathways (Cueva et al., 2020; Elinav et al., 2019; Tsvetikova and Koshel, 2020). Among metabolites, S-adenosyl methionine or SAM has emerged as the one that regulates the balance between cell survival and cell death. Produced by bacteria as well as by mammalian cells, SAM regulates cysteine-methionine metabolism, immune response and methylation of nucleotides thus controlling transcriptional processes. SAM has been used as an anti-tumor agent as well (Mahmood et al., 2020; Mehdi et al., 2020). Among metabolites solely made by bacteria is a t-RNA homolog, Queuosine (Q). Q cannot be synthesized by mammalian cells; however, it is required by mammalian cells for tRNA modifications (Morris and Elliott, 2001; Slany and Kersten, 1994; Tuorto et al., 2018). Originally identified in E. coli, queuosine was found to occupy the first anticodon position of tRNAs for histidine, aspartic acid, asparagine and tyrosine (Fergus et al., 2015). The hyper-modified nucleobase of queuosine is queuine. In mammalian cells, queuine treatment is reported to modulate tolerance to hypoxia (Reisser et al., 1994), influence proliferation (Langgut and Kersten, 1990; Langgut et al., 1993) and the expression of lactate dehydrogenase(Pathak and Vinayak, 2005).

Apart from altering gut microbiome, obesity also affects the therapy resistance by enriching for therapy resistant population in breast cancer (Bao et al., 2020) and ovarian cancer (Huang et al., 2020). Signaling between adipokines from adipocytes and cancer cells are responsible for this enrichment. Whether similar enrichment occurs in pancreatic tumors is not known. Recently published study from our laboratory show that pro-tumorigenic cytokine IL6 can enrich for therapy resistant CD133+ population in pancreatic cancer cells (Kesh et al., 2020a). Interestingly, we as well as others show that IL6 is the major cytokine produced under obese conditions (Hayashi et al., 2018; Wunderlich et al., 2018).

In this study, we observed that in high fat diet fed obese animals, pancreatic tumors did not respond to standard of care treatment like Gemcitabine/Paclitaxel cocktail. Interestingly, when the microbial composition of the lean and the obese animals was changed by fecal microbial transplant (FMT), the obese mice started responding to therapy, indicating a direct role played by the gut microbiome on therapy response. A deeper analysis of the microbiome revealed an enrichment of bacteria secreting the metabolite queuosine (Q) in the obese animals and an enrichment of S-adenosyl methionine (SAM) secreting bacteria in the lean animals. We further show that both high fat as well as treatment with Q upregulated PRDX1, an antioxidant protein that protects tumor cells from chemotherapy induced oxidative stress. In addition, high fat diet fed mice showed an enrichment of CD133+ treatment refractory cells that show high drug detoxification properties. Thus, the current study shows a potential dichotomous role of obesity in inducing therapy resistance: at the systemic level, obesity induced enrichment of Q producing bacteria that is protects tumors from oxidative stress and at the microenvironmental level, obesity enriches for therapy resistant population within the tumors.

## Material and Method

### Experimental animal

C57BL/6J mice were obtained from the Jackson Laboratory (Bar Harbor, Maine, USA). All mice were male and 4–6 weeks old. Food and tap water were available ad libitum. All mice were housed four mice per cages and maintained on a 12-h light/dark cycle, in a constant temperature (72⍰1±⍰1⍰°F) and 50% humidity. All procedures were conducted according to the protocols approved by the University of Miami Institutional Animal Care and Use Committee (IACUC).

### Animal model for diet induce obesity

Thirty-two male C57BL/6J mice were first randomly divided into 2 groups [Obese group, and Lean group]. Mice in the Obese group were given a high caloric diet brand name TD.88137, adjusted calories diet (42% from fat) and Lean group feed adjusted control diet named TD.08405 adjusted control diet (4% from fat) (Envigo, USA) during the total experiment period. After 4 weeks on designated diet, the weight gain was monitored for both groups. Additionally, blood glucose levels, serum triglyceride levels and blood cholesterol levels were measured to validate establishment of model.

### Blood glucose, triglyceride and cholesterol measurement

Blood samples were collected by retro-orbital sinus puncture via the medial canthus of the eye using clean 44.7-μ L heparinized micro hematocrit tubes. No anesthesia was used at the time of the blood sampling, to avoid unequal variations between animals and avoid the effects of anesthesia on the blood glucose levels. Mice blood glucose was measured using true track blood glucose meter and strips (Trivida health). Measurement of total cholesterol and triglyceride from mice serum were performed using cholesterol assay kit (Abcam) and triglyceride assay kit (Sigma-Aldrich) according to the manufacturer’s instructions.

### Tumor implantation

5×10^3^ number of pancreatic cancer cells (KPC001) were implanted in both groups of mice after 4 weeks of diet. After 2 weeks of tumor implantation when subcutaneous pancreatic cancer model was established, the 2 groups of mice were each further randomly divided into 2 subgroups (Obese and Obese + Gem/Pac; Lean and Lean +Gem/Pac). The Gem/Pac groups in both obese and lean mice received intraperitoneal injections of 100 mg/kg of gemcitabine and 10 mg/kg of paclitaxel twice in a week for consecutive 4 weeks, while the HF and LF groups were only receive equal volume of saline. In another set of experiment 32 mice were similarly divided in to 2 groups which further subdivided in 4 groups (Obese, Lean, Obese+Gem/Pac, Lean+ Gem/Pac) after 1 month feeding. These mice served as tumor free control and sacrifice at similar time point.

### Reciprocal Fecal microbial transplantation experiment

48 mice were divided in to two groups named Obese(donor) and Lean(donor) and received High Fat and Adjusted Control diet respectively. These animals serve as a fecal microbial donor pool. After one month of feeding animals were sacrificed and stool sample were collected aseptically. Stool were preserved at - 80C for future used. 200mg of the fecal extract was suspended in 1mL sterile PBS, filtered through 70µM cell strainer, and centrifuged at 6000Xg for 20min. About 10^10^ CFU/mL fecal bacteria were suspended in 6% NaHCO_3_ buffer with 20% sucrose. 32 mice were divided into Obese and Lean group (16 mice each) and feed for 1 month with High Fat and Adjusted Control diet. Obese mice were further randomized into Obese, Lean>>Obese(FMT) and Lean mice divided into Lean, Obese>>Lean(FMT). Lean>>Obese(FMT) mice were continuously feed in HF diet but received oral gavage of Lean mice stool for 3 time in a week for continuous 6 weeks similarly Obese>>Lean(FMT) received HF stool remain in adjusted control diet.

### In vivo treatment of SAM

64 mice were divided equally and feed with high fat and adjusted control diet separately for 1 month. HF group further dived in to 4 groups as Obese, Obese+GP, Obese+SAM, Obese+GP+SAM. Lean groups also divided similarly. Subcutaneous pancreatic tumor were implanted in each groups at day 30. From day 45 onwards treatment were started with either Gem/Pac alone or with SAM (Sigma Aldrich) or SAM alone. SAM were dissolve in saline and given 100mg/kg BW every day for 4 weeks. Mice were sacrificed. Tumor volume and weight were measured.

### Fecal matter collection and DNA Isolation

Fecal samples were collected in different time point to understand the effect of several sequential treatment in gut microbiota. Fecal samples collection was performed at day 1, 45, 60 and 90. Samples were collected in a sterile Eppendorf tube inside a biosafety cabinet with sterile forceps. Each group consisting eight animals were randomized (group wise) to nullify cage-effect in microbiome studies among the groups. After 90 days, all animals were sacrificed according to protocols approved by University of Miami Animal Care Committee. Part of the tumor sample were flash frozen in liquid nitrogen, while the rest were formalin fixed for paraffin embedding and histochemical analysis. Blood was collected by cardiac puncture prior to euthanizing the animals. Serum samples were stored for biochemical analysis. DNA from the murine fecal samples was isolated using the Power Soil DNA Isolation Kit (Qiagen) according to manufacturer’s instructions. All samples were quantified using the Qubit® Quant-iT dsDNA High-Sensitivity Kit (Invitrogen, Life Technologies, Grand Island, NY) to ensure that they met minimum concentration and mass of DNA and were submitted to University of Minnesota Genomics Center for Whole Genome Sequencing.

### Metagenomic Sequencing and Microbiome analysis

Shotgun metagenomic library was constructed from fecal DNA with the Nextera DNA sample preparation kit (Illumina, San Diego, CA), as per manufacturer’s specification. Barcoding indices were inserted using Nextera indexing kit (Illumina). Products were purified using Agencourt AMpure XP kit (Beckman Coulter, Brea, CA) and pooled for sequencing. Samples were sequenced using MiSeq reagent kit V2 (Illumina) in a HiSeq2500 sequencer.

Raw sequences were sorted using assigned barcodes and cleaned up before analysis (barcodes removed and sequences above a quality score, Q≥30 taken forward for analyses). For assembly and annotation of sequences, MetAMOS (Treangen et al., 2013) pipeline or Partek Flow software (Partek^®^ Flow^®^, Partek Inc., St. Louis, MO) were used. These softwares provide powerful tools to filter unique hits between human and mouse-specific genes versus microbial signatures. Alpha and Beta diversity calculations were done using embedded programs within the metagenomic pipeline, or using Stata15 (StataCorp LLC, College Station, TX) or EXPLICET software(Robertson et al., 2013). Functional profiling was performed using HUMAnN2-0.11.1 (Abubucker et al., 2012) with Uniref50 database to implement KEGG orthologies.

### Histology and and TUNEL assay

Tumor from all groups of mice were sectioned for histological studies. The tissue samples were fixed in 10% formalin and embedded in paraffin. The sections (5 μm) were cut using microtome, stained with hematoxylin and eosin, and slides were assessed using microscope (Leica microsystems, Germany) using at original magnification 10X and processed in Adobe Photoshop. For TUNEL study, paraffin embedded sections were deparaffinized with xylene followed by rehydration with descending alcohol series. Study was performed according to the manufacturer protocol (Abcam).

### Sirius red staining and measurements

Tissue sections were deparaffinized and hydrated in a descending order of alcohol solution, followed by PBS washing. Collagen staining were performed using picrosirius red staining solution (Chondrex Inc). The sections were washed with acidified water and dehydrated using absolute alcohol followed by mount in a resinous medium. The Sirius red–stained area was quantified using ImageJ software by selecting stained fibers in randomly selected five fields at a magnification of 10x under a light microscope.

### Cell culture

Pancreatic cancer cell line MIAPaCa-2 and SU86.86 was purchased from ATCC, KPC001 was isolated from the KRAS^G12D^ TP53^R172H^Pdx-Cre spontaneous mouse model from pancreatic cancer; S2VP10 was a gift from Masato Yamamoto (University of Minnesota, MN); colon cancer cell line (SW620 and RKO) were a gift from David Robbins (U Miami). S2VP10 was cultured in RPMI supplemented with 10% fetal bovine serum (FBS) with 1% Pen Strep (Life Technologies). All others were cultured in DMEM high glucose (Hyclone) containing 10% fetal bovine serum (FBS) with 1% Pen Strep (Gibco). All the established cell lines were used between passages 5 and 18. All cells were maintained at 37 °C in a humidified air atmosphere with 5% CO_2_. 70% confluent cells were used in each experiment. Cell lines were routinely checked for mycoplasma contamination and verified by STR profiling.

Small molecules inhibitor pre-queuosine 1 (50uM, Sigma Aldrich), SAM (200 uM, Sigma) and paclitaxel (50 nM, Sigma Aldrich) were used in various experiment.

### Isotope Ratio Outlier Analysis (IROA)

IROA was done on serum samples from animals on high fat and control diet following protocols described in (Mendez et al., 2019)and analyzed in Q-Exactive mass spectrometer.

### Statistical analysis

Data were presented as the Mean ± SEM. Statistical analyses were performed using GraphPad Prism software, version 8.0. Differences between two groups were analyzed by Student’s *t* test. *P* < 0.05 was considered statistically significant. Most statistical functions for microbiome and metabolome were embedded within MetAMOS (Treangen et al., 2013) pipeline or Partek Flow software (Partek^®^ Flow^®^, Partek Inc., St. Louis, MO). Output files from microbial sequence analysis and predictive metabolomics were further subjected to groupwise comparison. Depending on the analysis (as mentioned in respective figure legends), test of significance was either Mann-Whitney U test (Graphpad Prism), one-way ANOVA or 2-tailed t-test with False Discovery Rate (FDR) correction (using Bonferroni or Benjamini-Hochberg correction). The FDR threshold was set at 0.1 and p<0.05 was considered to be significant.

### Ethics statement

All animal studies were performed according to the protocols approved by IACUC at University of Miami, USA in accordance with the principles of the Declaration of Helsinki. All authors had access to all data and have reviewed and approved the final manuscript.

## Result

### Diet Induced hyperlipidemia deregulated gut microbiome in the tumor bearing obese mice

To determine the change in the microbiome, we analyzed the fecal microbiome in lean and obese animals after the animals were put on high fat diet and (and adjusted control diet) for 30 days according to the schema **Figure 1A**. Collection L1-L4 and O1-O4 represented fecal matter collection at different stages: L1/O1: Start of experiment; L2/O2: Effect of diet on microbiome; L3/O3: Effect of implanted tumor and its growth in lean and obese mice; L4/O4: End point collection of fecal matter from animals that did not receive Gem/Pac and L4’/O4’: Effect of Gem Pac therapy on microbiome change in lean and obese animals. Our data showed an expected change in the microbiome between the control (lean) and high fat fed (obese) mice (**Figure 1B**). Since each animal was tracked throughout the experiment, we next determined the changes in the composition of microbiome as a consequence of diet in a temporal fashion. Analysis of O1-O4 collection showed visible changes in the microbiome compared to L1-L4 indicating the diet induced hyperlipidemia and obesity changed the microbiome both before and after tumor implantation (**Figure 1C**). At the phylum level, there was slightly increased abundance in Proteobacteria and Firmicutes in animals on high fat diet for 30 days (Collection O2 compared to collection L2). Similarly, we observed a decrease in the Bacteroidetes in these animals (**Figure 1D**). This relative abundance was maintained through collection O3 and O4 (**Figure 1E**). Similar relative abundance was observed at the class level as well (**Supplementary Figure 1A**). When we compared the gut microbiome in response to Gem-Pac treatment within each group, there was no observable difference in the relative abundance between the lean and obese animals treated with Gem/Pac (**Supplementary Figure 1B)**.

**Figure 1.**
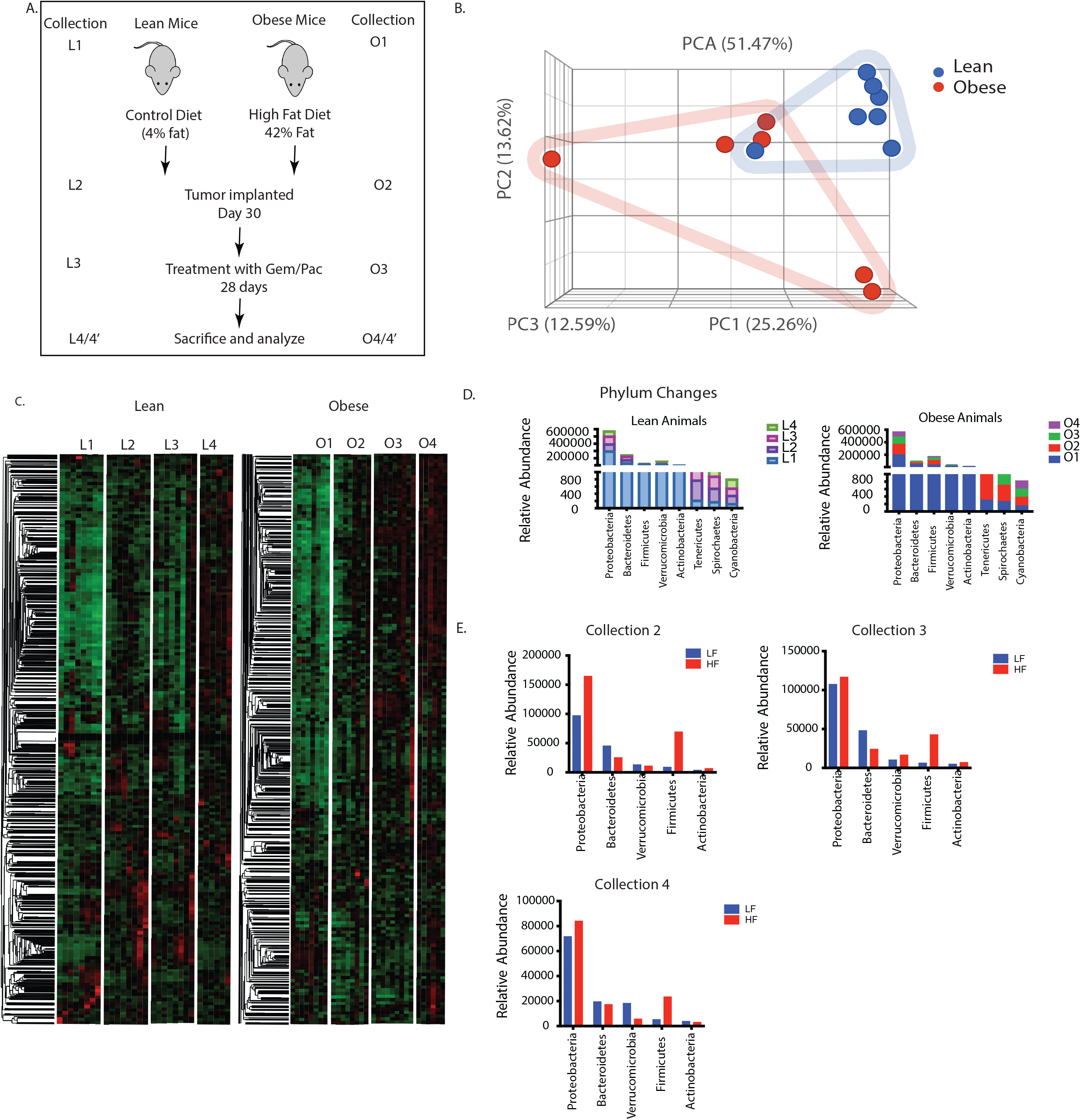
Obesity induced change in gut microbiome: Schematic diagram showing timeline of the experiment (A). High fat diet changed the gut microbiome in C57Bl6 mice (B). Differences in the microbial composition can be visualized in the heat map L1= 1^st^ collection in lean mice before tumor implantation; L2= 2^nd^ collection in lean mouse after 1 month of diet; L3=3^rd^ collection before start of Gem/Pac therapy in lean mouse, L4= Final collection after sacrificing in lean mouse. O1=1^st^ collection in obese mice before tumor implantation; O2= 2^nd^ collection in obese mouse after 1 month of diet; O3=3^rd^ collection before start of Gem/Pac therapy in obese mouse, O4= Final collection after sacrificing in obese mouse (C). Dynamic changes in gut microbiome in phylum over the collection period (D,E).

### Obesity mediated fecal microbiome confer resistance to standard of care in PDAC

Animals on high fat diet predictably showed increase in body weight after 30 days on their diet, whether adjusted control/lean or high fat/obesogenic (**Supplementary Figure 2A**). Similarly, an increase in blood glucose, triglyceride and cholesterol levels was observed in the obese mice on high fat diet compared to those on adjusted control diet (**Supplementary Figure 2B-D**). There was no significant difference in the tumor take between the two groups (**Supplementary Figure 2E**). In the animals on high fat diet, the tumor progressed faster compared to those in the adjusted control diet (**Supplementary Figure 2F**) and showed accumulation of lipids (**Supplementary Figure 2G**). Endpoint observation showed that animals in the lean group (on an adjusted control diet) responded to Gem-Pac regimen (**Figure 2A, B**) while those on the high fat diet failed to respond (**Figure 2 C,D**). Interestingly, upon changing the microbiome composition of the two groups of animals (lean and obese) by transplanting the fecal microbiome of lean mice to obese and vice versa right before tumor implantation (As shown in Schema **Figure 2E** and PCA plots **Figure 2F, G**), we observed that the response of the tumors was reversed. Lean animals that previously responded to chemotherapy, stopped responding to Gem/Pac after receiving obese FMT (**Figure 2H**) and obese animals that we resistant to Gem/Pac started responding to this therapy after receiving the lean FMT (**Figure 2I**). H&E analysis of the tumor samples from the groups showed expected accumulation of infiltration of fat droplets in the tumor tissue in the obese mice compared to the lean mice. Further, obese mice receiving lean FMT showed extensive areas of necrosis in the histology (**Figure 2J**). Similarly, these obese mice (that received lean FMT also showed widespread fibrosis and collagen deposition that was not observed in the other groups (**Figure 2K**). Additionally, this group of obese animals receiving lean FMT also had large areas of apoptotic cells (**Figure 2L**). This indicated that the gut microbiome played a role in driving therapy resistance in obese animals.

**Figure 2.**
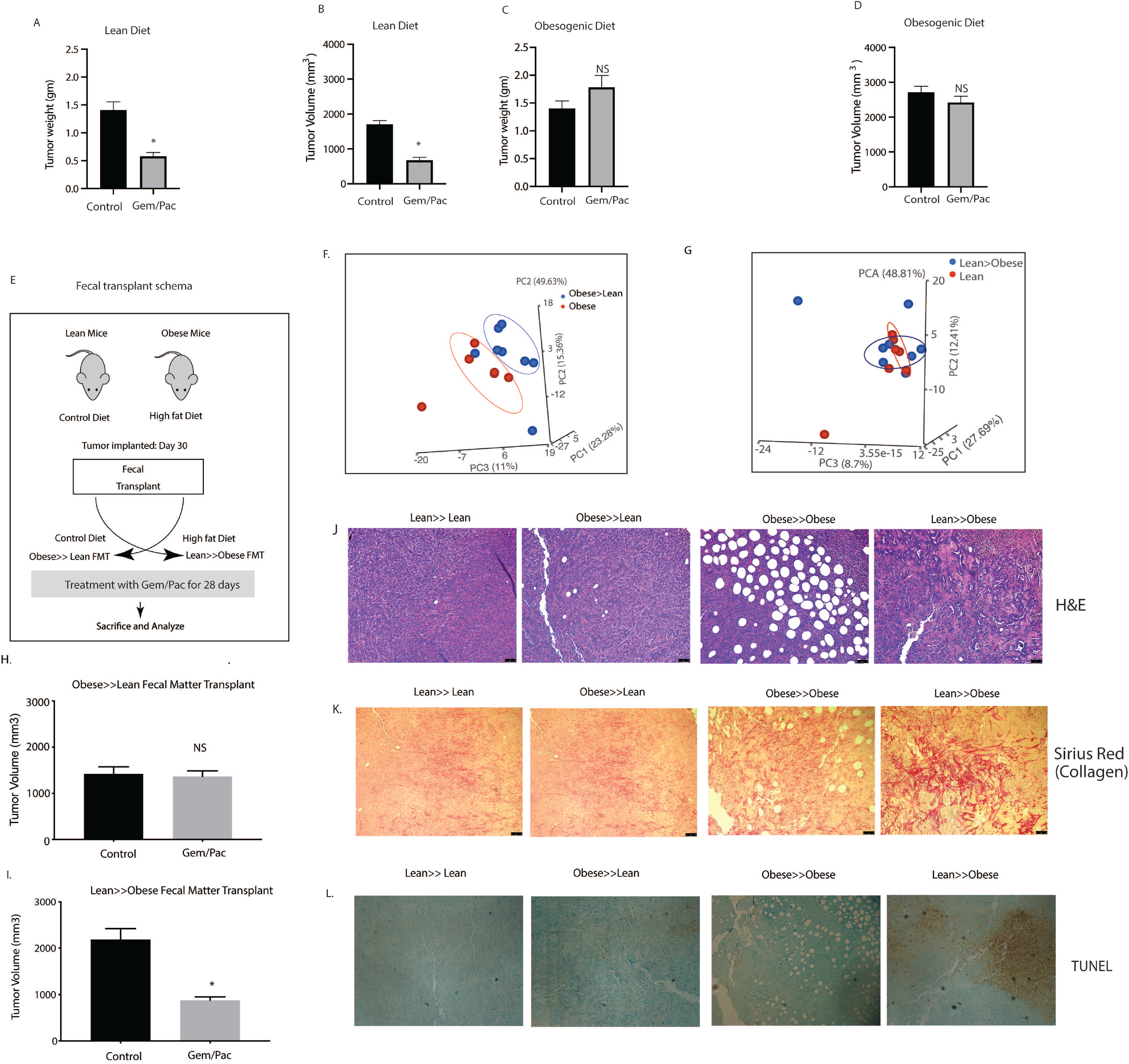
High fat diet fed mice show resistance to chemotherapy. KPC001 cells were implanted subcutaneously in C57BL6 mice on control and high fat diet and treated with Gem/Pac for 4 weeks. Tumor weight (A) and tumor volume (B) in mice on control (lean) diet showed significant decrease. Tumor weight (C) and tumor volume (D) in mice on high fat (obesogenic) diet showed no significant response. Schema for fecal transplant (E) is shown in which the high fat diet fed mice received the lean mice microbiome and vice versa. PCoA plots of Obese to lean transplant (F) and lean to obese transplant (G). Obese>>Lean FMT showed loss of response in the presence of chemotherapy (H) while Lean>>Obese FMT showed sensitivity to chemotherapy (I). Visible changes in histology was observed in the Lean>>Obese FMT in H&E slides (J), collagen deposition (K) and TUNEL staining (L).

### Diet induced hyperlipidemia enriched for tumor-protective microbial metabolic pathways

Since the change in gut microbiome due to high fat diet contributed to resistance to Gem/Pac therapy, we next used the WGS of the gut bacteria in lean and obese mice to study the enrichment of microbial population that might correspond to this phenomenon. Our analysis showed that a distinct microbial population was enriched in the obese mice when compared with the lean. These included bacteria that were enriched for biosynthesis of queuosine, a tRNA homolog, contributing to protection of oncogenesis related stress (**Figure 3A,B**). Similarly, the analysis of the lean mice microbiome, showed an enrichment of bacteria metabolizing S-adenosylmethionine or SAM (**Figure 3C,D**).

**Figure 3.**
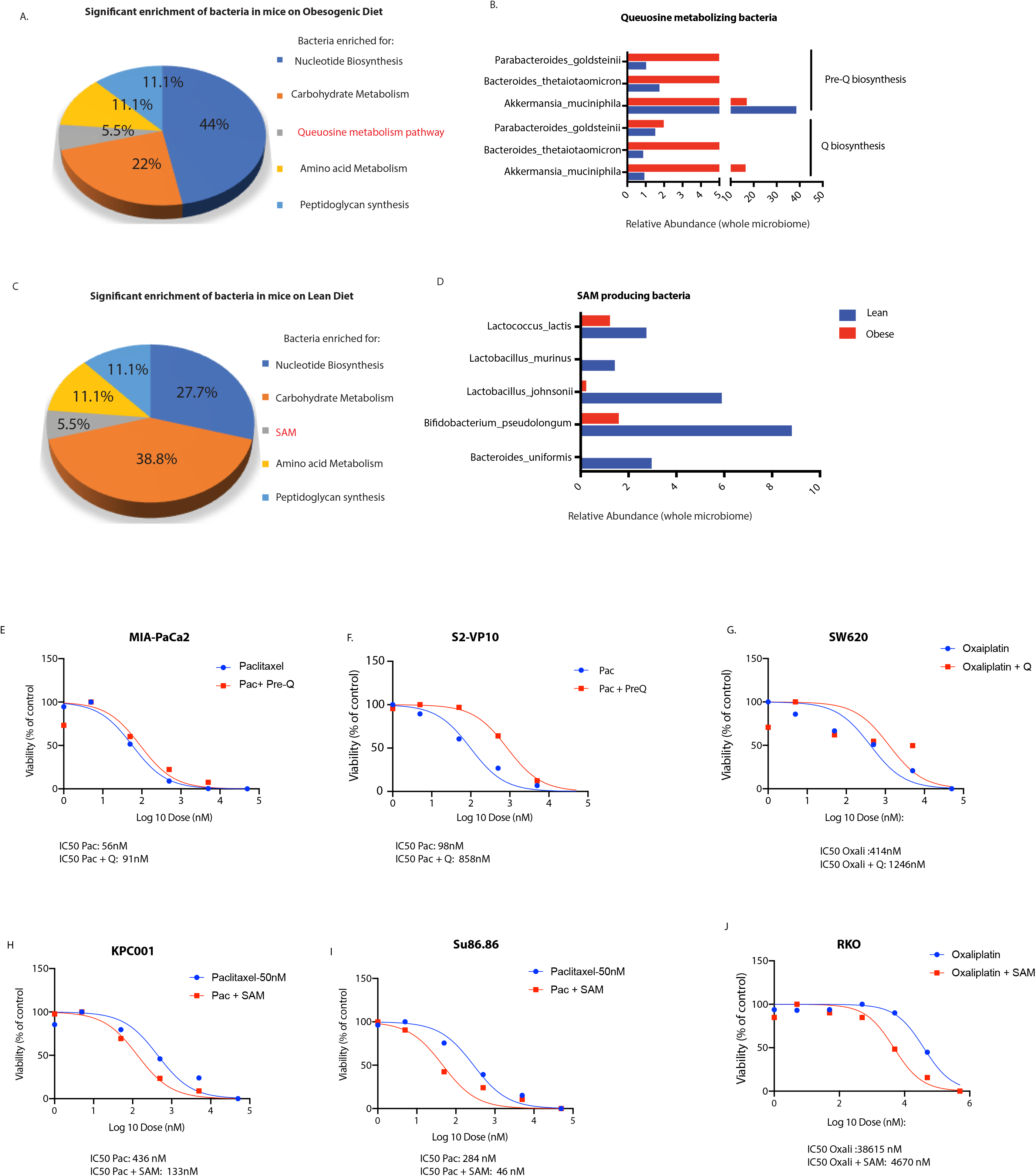
Metabolomic reconstruction using humaN2 pipeline was performed to determine the microbial metabolome. High fat diet fed mice showed an enrichment of Q metabolizing bacteria (A, B) while lean mice showed an enrichment of SAM metabolizing bacteria (C,D). Treatment of pancreatic cancer cells MIA-PACA2 (C) and S2VP10 (D) with paclitaxel and pre-Q showed a shift in IC50 indicating resistance. Treatment of colon cancer cell SW620 with oxaliplatin and pre-Q showed a similar shift in IC50 (E). Treatment of pancreatic cancer cells KPC001 (F) and Su86.86 (G) with SAM showed an opposite shift of IC50 indicating sensitization. Treatment of colon cancer cell RKO (H) with oxaliplatin and pre-Q showed a similar shift in IC50 indicating sensitization by SAM.

To demonstrate that treatment with queuosine protects tumor cells from drug induced stress, we next treated paclitaxel sensitive MIA-PACA2 and S2VP10 cells with queuosine precursor (Pre-Q) since queuosine cannot be taken up by the cells. Our studies showed that treatment with pre-Q increased proliferation of both cancer cell lines (**Supplementary Figure 3 A, B**) and protected them from paclitaxel induced cell death (**Figure 3E, F**). Similar rescue was also observed in SW620 colon cancer cells (**Figure 3G)**.

Since our hypothesis was that microbial metabolites from lean mice sensitized the tumors to chemotherapy, we next validated if treatment with SAM sensitized normally resistant pancreatic cancer cells to Gemcitabine and Paclitaxel. We treated paclitaxel resistant SU86.86 and KPC001 cells with indicated doses of SAM along with paclitaxel and observed that SAM reverted paclitaxel resistance in these cells (**Figure 3H, I**). Similar reversal of resistance was also observed in oxaliplatin resistant colon cancer cell line RKO (**Figure 3J**). The changes in IC50 with Q and SAM in the presence of chemotherapy compound is shown as part of Figure 3J.

### Serum metabolites of obese animals showed enrichment of nitrogen metabolism and detoxification pathways

To confirm if these pathways were enriched in the serum samples as well, we next performed a Isotypic Ratio Outlier Analysis (IROA). A total of 217 different metabolites were detected in the serum of the lean and obese animals. Complete list of metabolites identified are included in Supplementary Table 1. A Metabolite Set Enrichment Analysis (MSEA) with the identified metabolites showed a significant enrichment of metabolites in aminoacyl tRNA biosynthesis followed by those involved in phenylalanine, tryptophan and tyrosine metabolism **(Figure 4A)**. Upon separating the identified metabolites into nodes, we observed that the major node involved nitrogen metabolism encompassing urea cycle, ammonia recycling and glutamine/glutamate metabolism. Similarly, another distinct metabolic node identified was that of the mitochondrial oxidation cycle that contributes to ROS accumulation and beta-oxidation **(Figure 4B)**. Upon comparing the relative abundance of the metabolites identified by IROA, L-glutamic acid/glutamine metabolism (that feeds into glutathione, proline and arginine metabolism, ammonia recycling as well as urea cycle), was found to be the most abundant metabolite in obese animals **(Figure 4C, D)**. An in-depth analysis of the metabolic pathways identified was next performed in MetaCyc database (Caspi et al., 2020). We observed that there was a general upregulation of biosynthesis, degradation and energy pathways in obese animals (**Figure 4E**). Further, the obese animals tended to have increased accumulation of metabolites in the detoxification process, specifically glutathione (**Figure 4F**). This was consistent with the increased glutamic acid/glutamine accumulation in obese mice.

**Figure 4.**
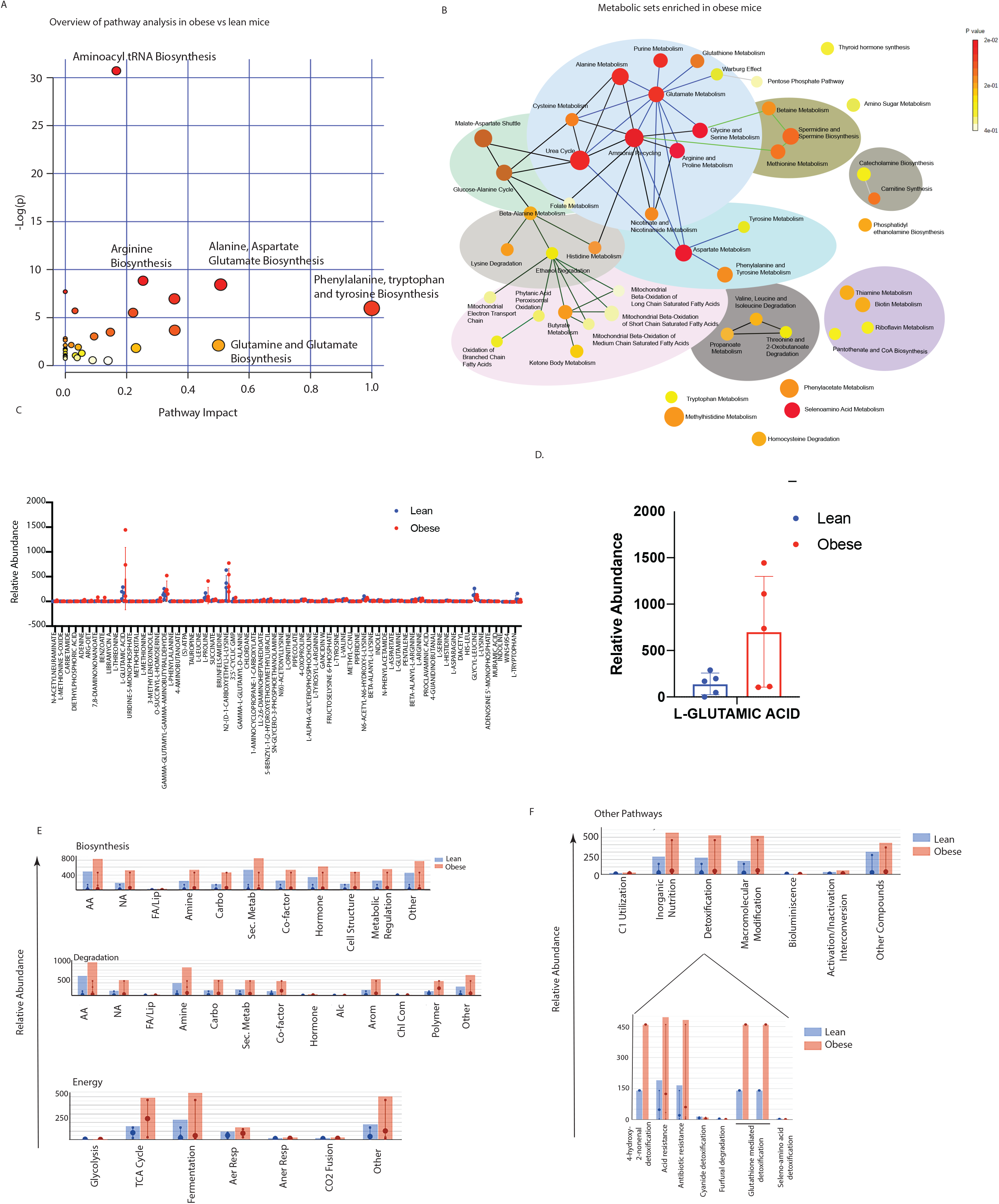
Serum metabolomics of lean and obese animals using IROA: Pathway Analysis of obese vs lean animals using MetaboAnalyst showing enrichment of critical pathways (A). Metabolic Set Enrichment analysis (MSEA) of obese animals (B). Relative abundance of metabolites in lean vs obese animals (C). Obese mice showed an increase in glutamic acid (D). Metacyc showed relative changes in the metabolic pathways in lean and obese animals (E). Drug detoxification pathways were enriched in obese animals (F).

### S-Adenosylmethionine (SAM) is present as a fecal metabolite and sensitizes pancreatic tumor cells to chemotherapeutic agents

Since our metabolomic reconstruction of the WGS of the gut bacteria in the lean animals showed an enrichment of SAM metabolizing bacteria and we did not detect SAM in the serum, we next estimated SAM from the fecal samples of the lean and obese mice. Our ELISA based analysis showed that SAM was significantly elevated in the lean mice (**Figure 5A**). Obese mice did not have abundant SAM in their fecal samples but upon lean microbial transplant, they started producing SAM (**Figure 5A**). Similarly, tumors from lean mice showed increased accumulation of SAM in lean animals compared to the tumors in obese animals (**Figure 5B**). To study if SAM sensitized pancreatic tumors to Gem/Pac therapy in vivo, we then implanted tumors in lean and obese mice as demonstrated in **Figure 5C**, and treated the animals with SAM in the presence of Gem/Pac standard of care. Tumor bearing obese mice that received SAM showed greater sensitivity to Gem/Pac regimen compared to those that did not receive SAM. There was significant reduction in tumor volume (**Figure 5D**) and weight (**Figure 5E**). Tissue histology showed decreased fibrosis as observed by Sirius red staining (**Figure 5F**) and less Ki67+ cells (**Figure 5G**).

**Figure 5.**
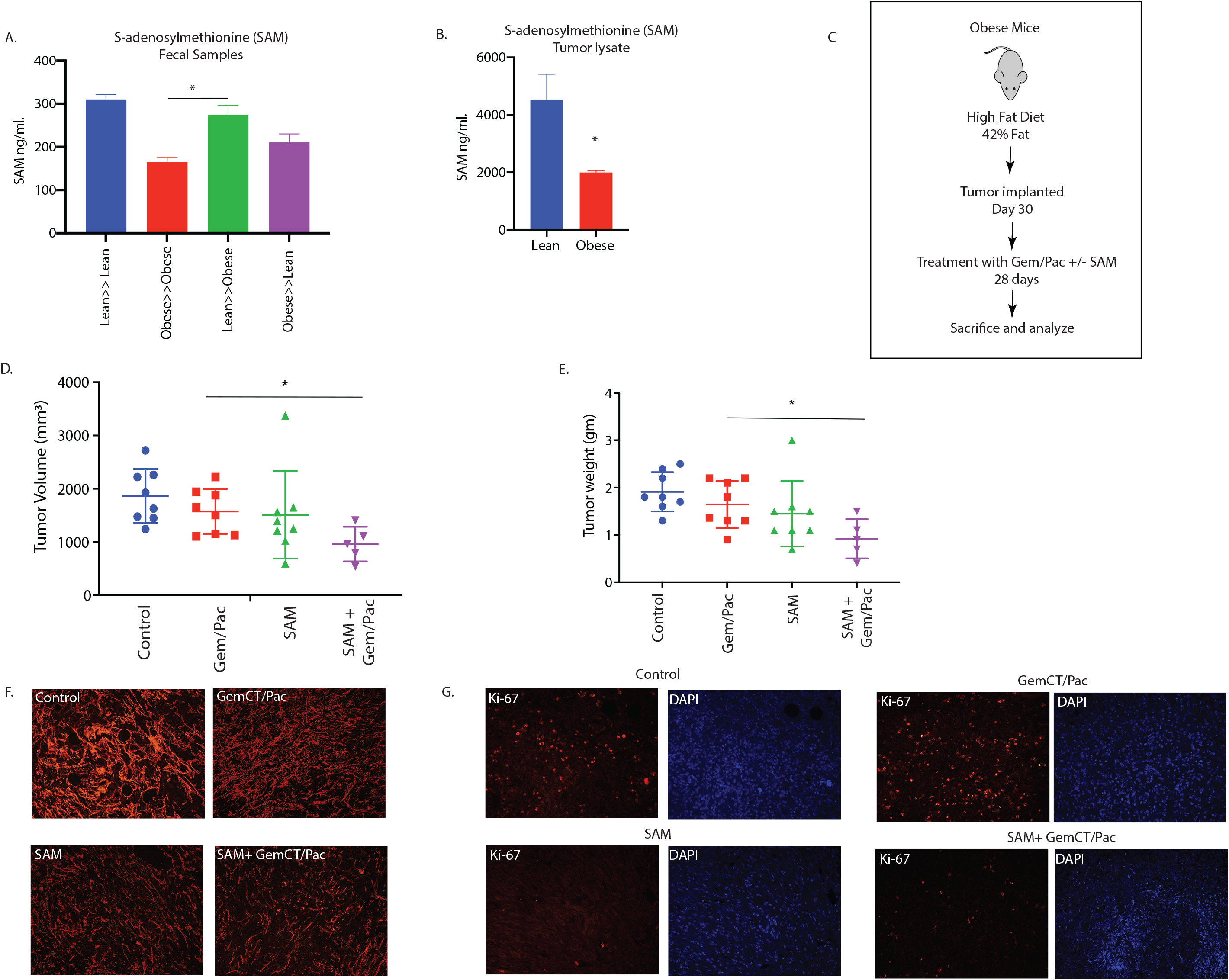
SAM reverted effect of obesity induced therapy resistance: SAM was increased in Lean>>Obese FMT animals in fecal (A) sample. SAM was decreased in Obese animals (B). Schema showing experimental set up SAM treatment in high fat diet fed, pancreatic tumor bearing mice (C). SAM decreased tumor volume (D) and weight (E). Treatment with SAM also decreased collagen (F) and Ki67+ cells (G).

### Queuosine induces PRDX1 expression in pancreatic cancer cells to promote resistance to chemotherapy

Chemotherapy compounds induce oxidative stress and generate reactive oxygen species (ROS) in cancer cells resulting in cell death. Since Q has been implicated in antioxidant defense in cells (Pathak et al., 2008), we next studied the expression of genes involved in oxidative stress in the presence of Q using an oxidative stress PCR array. Our results showed that Q preferentially induced the expression of PRDX1 in pancreatic cancer cells (**Figure 6A**; **Supplementary Figure 4A-C**). Similar increase in Prdx1 protein expression was observed as well (**Figure 6B**). To study if chemoresistance in the presence of Q was via induction of PRDX1, we next inhibited PRDX1 using siRNA in pancreatic cancer cells SU86.86 and determined effect on cell viability in the presence of paclitaxel +/-Q. Silencing was verified by qPCR (**Supplementary Figure 4D**). Silencing PRDX1 did not alter the viability of the pancreatic cancer cells **(Supplementary Figure 4E)**. We observed that upon silencing Prdx1 in pancreatic cancer cells (MIA-PACA2), queuosine was unable to protect the cells from paclitaxel induced cell death **(Figure 6C)**.

**Figure 6.**
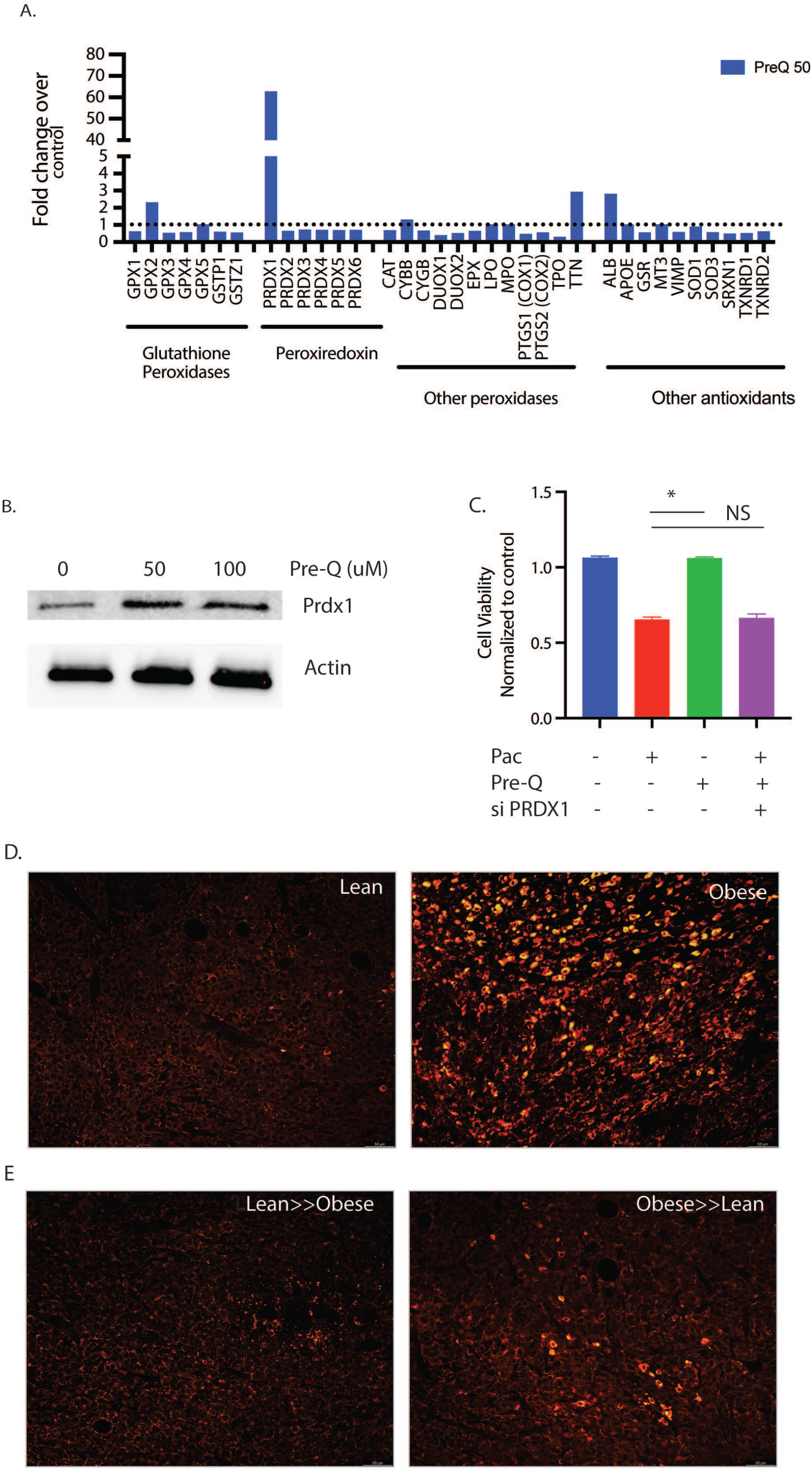
Queuosine mediated chemoresistance by upregulating PRDX1: Treatment of pancreatic cancer cell SU86.86 increased expression of PRDX1 as seen in oxidative stress PCR array analysis (A).Western blot showing upregulation of PRDX1 protein after treatment with Pre-Q (B). Silencing PRDX1 using siRNA presented Pre-Q induced resistance in MIA-PACA2 cells (C). IHC of lean and obese tumor bearing mice show upregulation of PRDX1 in obese mice (D). FMT of obese>>lean mice increased PRDX1 expression in these animals.

To confirm that in high fat fed mice had increased PRDX1 expression, we performed immunofluorescence on the tumors. Tumor bearing lean mice had low expression of Prdx1, while the high fat diet fed obese mice had a high expression of Prdx1 (**Figure 6D**). Upon fecal matter transplant from lean to obese decreased Prdx1 expression in obese mice while obese to lean fecal transplant increased the expression of this gene (**Figure 6E**), consistent with the chemoresistance observed in Figure 1.

### Obesity enriches for intra-tumoral treatment refractory population in pancreatic cancer

Obesity and adipocyte mediated signaling affects intra-tumoral oncogenic signaling along with causing system-wide inflammatory responses (Iyengar et al., 2016). Complete analysis of the serum cytokines using multiplexed assays showed increased IL6 among the elevated cytokines in the obese mice (Supplementary Figure 5A). Interestingly, our recently published research (Kesh et al., 2020a) showed that presence of IL6 enriched for a CD133+ treatment refractory population of cells in pancreatic cancer. Previous studies from our laboratory show that resistance to therapy in pancreatic tumors correlated with presence of CD133+ cells (Gupta et al., 2018; Nomura et al., 2016). Thus to study if obesity induced peritumoral adipocytes resulted in enrichment of CD133+ population in the tumor tissues, we next evaluated CD133+ cells in the tumors from obese mice. Our results showed that CD133+ cells were increased in these animals (compared to the lean animals) and tended to accumulate around the perilipin stained adipocytes in our animal models (**Figure 7A, Supplementary Figure 5B**). To study if adipocytes enriched for CD133+ therapy resistant population in pancreatic tumors, we next treated MIA-PACA2 (pancreatic cancer cells with negligible CD133+ population) with conditioned media from patient derived adipocytes. Our studies showed that treatment with adipocyte conditioned media made MIA-PACA2 cells resistant to paclitaxel (**Figure 7B**). Further analysis showed that this treatment also led to a distinct enrichment of CD133+ population (**Figure 7C, Supplementary Figure 5C**). Additionally, adipocyte conditioned media also enriched for stemness genes in the MIA-PACA2 cells (**Figure 7D-F**) Analysis of the adipocyte conditioned media also revealed IL6 as the major cytokine produced by the cells (**Supplementary figure 5D**). To study if queuosine increased stemness to promote chemoresistance, we next treated pancreatic cancer cells MIA-PACA2 and Su86.86 with pre-Q. We observed no change in the mRNA expression of stemness or self-renewal genes (**Supplementary Figure 5E**).

**Figure 7.**
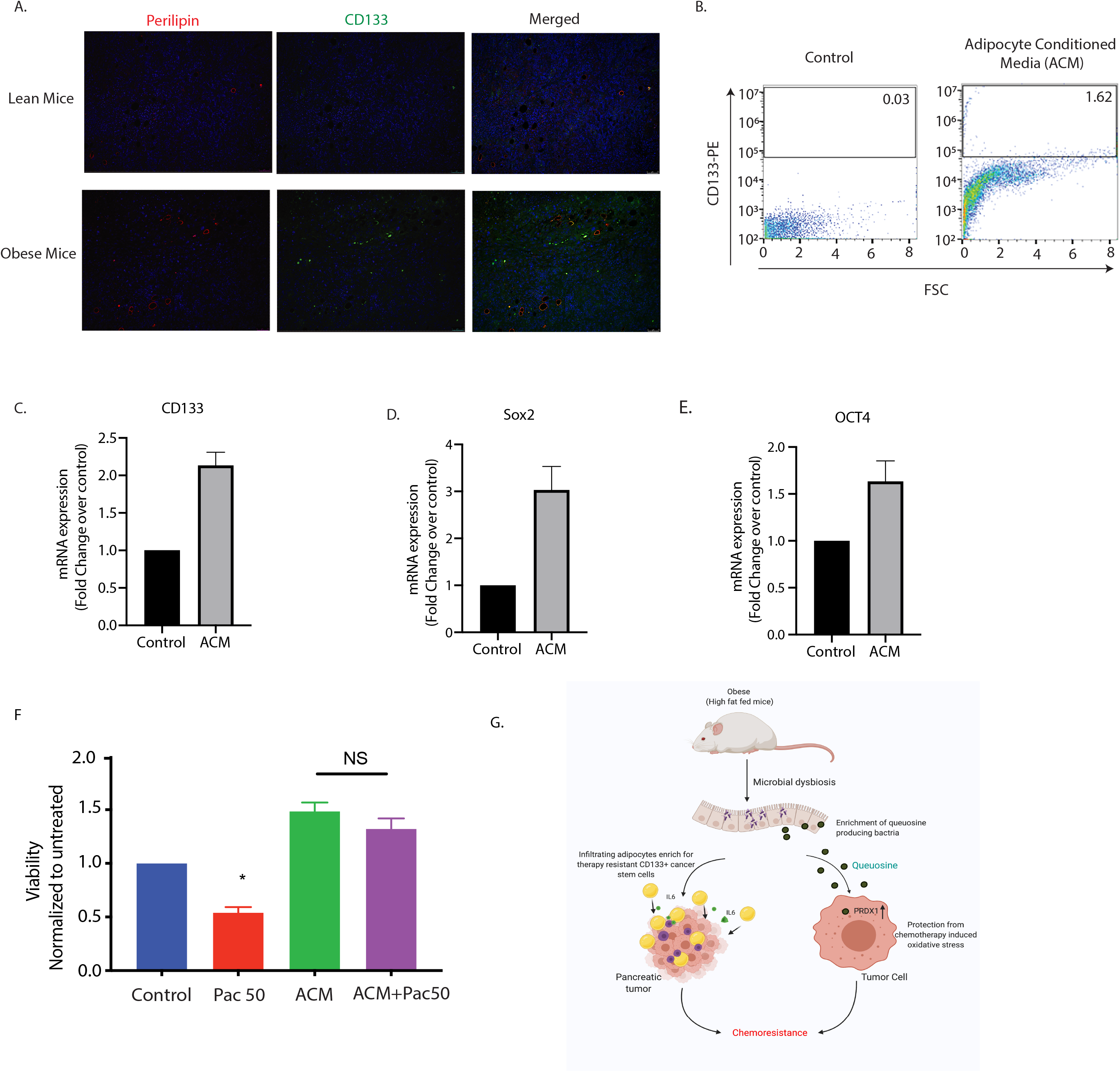
Obesity enriched for resistant CD133+ cells in pancreatic tumors. High fat diet fed tumor bearing mice showed an increase in CD133+ cells near lipid droplets (A). CD133+ population was increased when pancreatic cancer cells were treated with adipocyte conditioned media (B). Adipocyte conditioned media also increased expression of self-renewal genes like CD133 (C), Sox2 (D), Oct4 (E) and induced resistance to paclitaxel (F). Schematic diagram showing mechanism of obesity induced resistance.

## Discussion

Research on effect of obesity on different cancers has been gaining importance ever since onset of obesity at early years and sedentary lifestyle has become predominant. While systemic factors like chronic inflammation, hormones, circulating adipokines, and adipocyte-mediated inflammatory and immunosuppressive microenvironment and gut microbial dysbiosis have been attributed to the association of obesity with pancreatic cancer (Cani and Jordan, 2018; Cascetta et al., 2018; Pothuraju et al., 2018), adipocyte mediated intratumor processes have also been ascribed to poor prognosis of the disease. Over the past decade, growing evidence has shown that the composition of the gut microbiota and its activity might be associated not only with the onset of inflammation but also with metabolic disorders and cancer.

In accordance with this, studies from our laboratory have shown that gut microbiome is associated with pancreatic cancer progression (Mendez et al., 2020). Microbial dysbiosis has been correlated with therapy response in a number of cancers (Mima et al., 2017; Routy et al., 2018; Zheng et al., 2019). Our ongoing and recently published studies also indicated that conditions of high blood glucose (as seen in Type 2 diabetes) also contribute to chemoresistance in pancreatic cancer pre-clinical models (Kesh et al., 2020b). Similarly, tumor microbiota has been shown to contribute to gemcitabine resistance in pancreatic cancer (Geller et al., 2017). While the association of microbiome in tumor progression as well as in determining its properties are being evaluated, there is a lacuna in understanding the mechanism by which bacteria may affect these processes.

In the gut, the microbiome, its metabolites are in a constant state of flux with the host tissue and its secretome. Several microbial metabolites like short chain fatty acids (SCFA), trimethylamine (TMA) and polyamines have been directly implicated in driving tumorigenesis by contributing to protein and nucleotide synthesis for the rapidly proliferating tumor cells. Several studies show an enrichment of bacterial species that metabolize antioxidants and thus contribute to therapy resistance in multiple cancers (Bae et al., 2014; Kesh et al., 2020b; Oellgaard et al., 2017). In this study, we observed that in obese animals, pancreatic tumors did not respond to standard of care treatments like Gemcitabine/Paclitaxel cocktail. Interestingly, when the microbial composition of the lean and the obese animals was changed by FMT, the obese mice started responding to therapy. A deeper analysis of the microbiome revealed an enrichment of bacteria secreting the bacterial metabolite queuosine (Q) in the obese animals and an enrichment of S-adenosyl methionine (SAM) secreting bacteria in the lean animals. Q is a tRNA homolog, that protects from oncogenesis induced stress during tumor progression. Queuosine is also a modified nucleoside, the occurrence of which is widespread across the animal and plant kingdoms. Yet eukaryotes are unable to synthesize Q-nucleoside or any of its precursor forms. Instead, they salvage the nucleobase of queuosine, referred to as queuine or Q-base. In the case of metazoans, the source of queuine is dietary, whether from the gut microflora or from ingested food (Nishimura, 1983). In HeLa cells cultured in medium containing 10% horse serum, queuine treatment increased cell density under aerobic conditions but decreased cell density under hypoxic conditions (Langgut et al., 1990), suggesting that that queuine is a stimulant for proliferation in an aerobic environment, but inhibitory when conditions are hypoxic. Later, a study from the same group on the proliferation of non-transformed, transformed and tumor-derived cell lines concluded that queuine can stimulate or inhibit growth, depending on the cell line investigated (Langgut et al., 1993). Recent studies have shown the queuine can promote anti-oxidant defense system by activating cellular antioxidant enzyme activity in cancer (Pathak et al., 2008). It is well known that obesity mediated inflammation leads to generation of reactive oxygen species (ROS) in the cells. Interestingly, a study published in 2012 reported the structural organization of the enzyme GluQ-RS in bacteria, that was responsible for formation of GluQ tRNA modification(Caballero et al., 2012). This study and two others also showed that this enzyme required a high concentration of glutamate to be activated in the host so that it could be transferred to the queuosine base present on the tRNA^Asp^ (Dubois et al., 2004; Salazar et al., 2004). Thus, enrichment of queuosine metabolizing bacteria as well as accumulation of metabolites that promoted detoxification of xenobiotics in the tumor-bearing obese mice was indicative of protection from drug induced stress. Our study showed that treatment with pre-Q upregulated expression of PRDX1 in cells (Figure 6A,B) and obesity triggered its upregulation in tumor bearing animals (Figure 6D). Also, since silencing PRDX1 sensitized the pancreatic cancer cells to paclitaxel even in the presence of Pre-Q indicated that Q induced chemoresistance was being mediated via PRDX1 (Figure 6C).

SAM is known to be an anti-tumor metabolite. In gastric and colon cancer, SAM reverses hypomethylation status of c-myc and H-ras to inhibit tumor growth (Luo et al., 2010). Similarly, in breast cancer, treatment with SAM and doxorubicin showed anti-proliferative as well enhanced apoptotic properties (Ilisso et al., 2015). We detected SAM in the fecal as well as tumor samples of the tumor bearing lean animals. Based on this observation, we hypothesized that microbial metabolite Q accumulates in the obese animals and offers them protection from oxidative stress associated with chemotherapy. Upon transplanting the microbiome of the obese animals with that of the lean animals, this protection is lost, and the tumors start responding to chemotherapy. Similarly, accumulation of SAM in lean animals sensitizes the tumors in them to chemotherapy. When replaced with the obese microbiome, this sensitization is lost and tumors stop responding to chemotherapy. In this study we have validated this hypothesis and observed that treatment with pre-queuosine offers therapy resistance to pancreatic cancer cells, while treatment with SAM sensitizes them both in vitro and in vivo. Cancer stem cells are known to be enriched under conditions of obesity. Our data corroborated that (Figure 7). Adipocyte conditioned media showed enrichment in CD133+ population as well as increased expression of stemness genes. Interestingly, treatment with Q did not seem to affect the cancer stemness (Supplementary Figure 5E). This indicated that obesity induced poor prognosis and therapy resistance was being mediated both by enrichment of cancer stem cells from the pro-inflammatory cytokines secreted by the accumulated adipocytes in the microenvironment as well as by the microbial metabolite Q. This mechanism of resistance is summarized in Figure 7G.

## Conclusion

This study shows for the first time that microbial metabolites like Queuosine contribute to therapy resistance in pancreatic cancer under conditions of obesity by upregulating PRDX1, which protects them from chemotherapy induced oxidative stress. We further show that this therapy resistance can be reversed by FMT from lean mice as well as by SAM, another metabolite produced by gut bacteria. This finding can be of potential significance in case of pancreatic cancer patients that do not respond to standard of care as these “non-responding” patients can be made to “respond” to therapy by supplementing with S-adenosine methionine or SAM. This has immense potential in improving the survival statistics of patients diagnosed with pancreatic cancer.

## Acknowledgement

The authors would like to thank Dr. Bonnie Blomberg and Dr. Daniela Fresca for providing the adipocyte conditioned media, Dr. David Robbins for the colorectal cancer cell lines, Oliver Umland and University of Miami Flow cytometry core for the flow cytometry analysis.

## Funding

This study was funded by NIH grant R01-CA184274 and R01-CA124723 (to SB) and CA161976 (to NM); James Esther and King Biomedical Research Program by Florida Department of Health grant 9JK09 (to SB); pancreatic cancer action network grant to NM. Research reported in this publication was supported by the National Cancer Institute of the National Institutes of Health under Award Number P30CA240139.

## Conflict of Interest

University of Minnesota has a patent for Minnelide, which has been licensed to Minneamrita Therapeutics, LLC. SB is a consultant with Minneamrita Therapeutics LLC and this relationship is managed by University of Miami. The remaining authors declare no conflict of interest.

## Disclaimer

This article was prepared while Santanu Banerjee, PhD was employed University of Miami. The opinions expressed in this article are the author’s own and do not reflect the view of the National Institutes of Health, the Department of Health and Human Services, or the United States government.

