## Supplementary figures and images for "Diet induced hyperlipidemia confers resistance to standard therapy in pancreatic cancer by selecting for “tumor protective” microbial metabolites and treatment refractory cells"

### Supplementary Figures

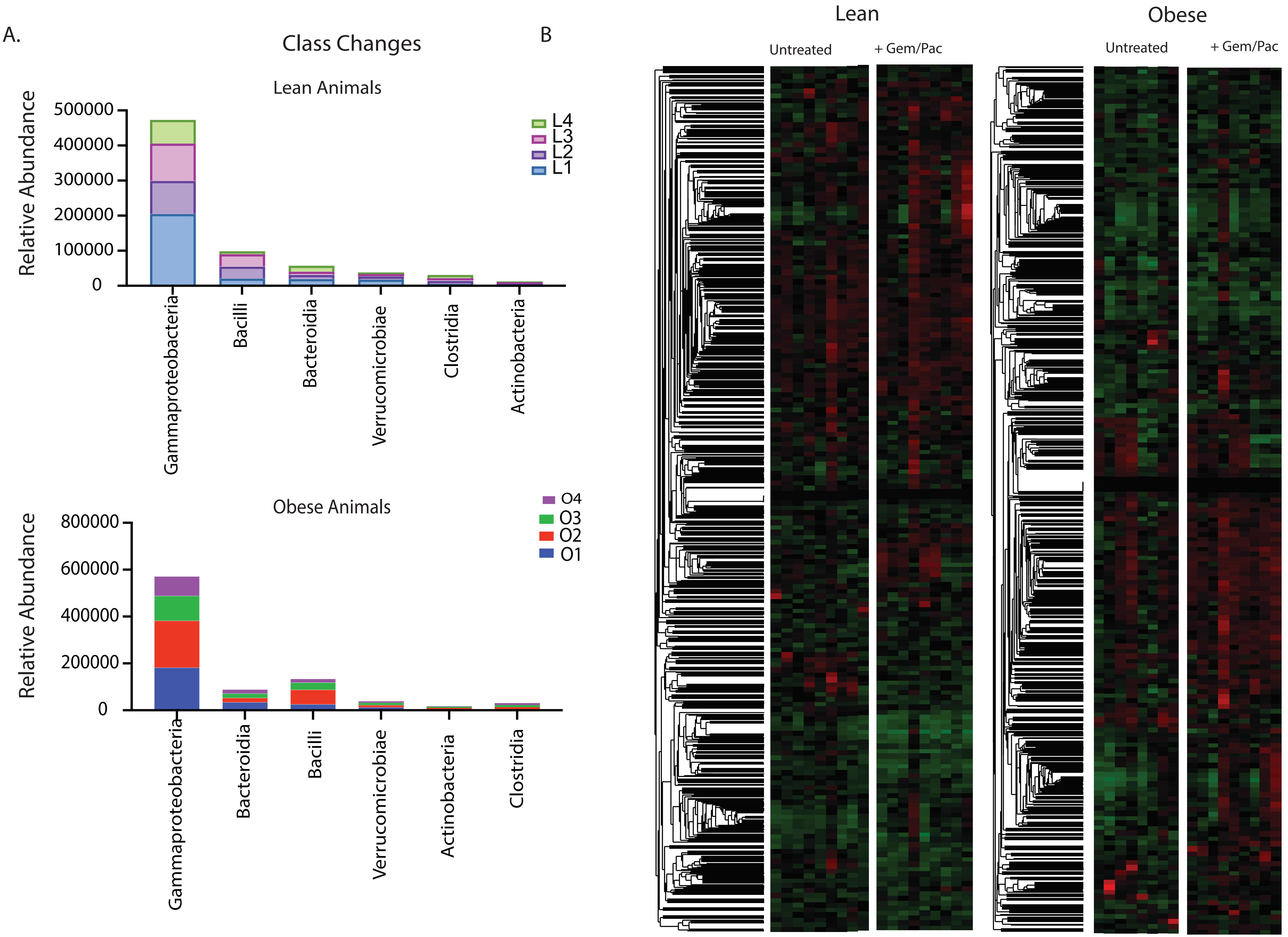

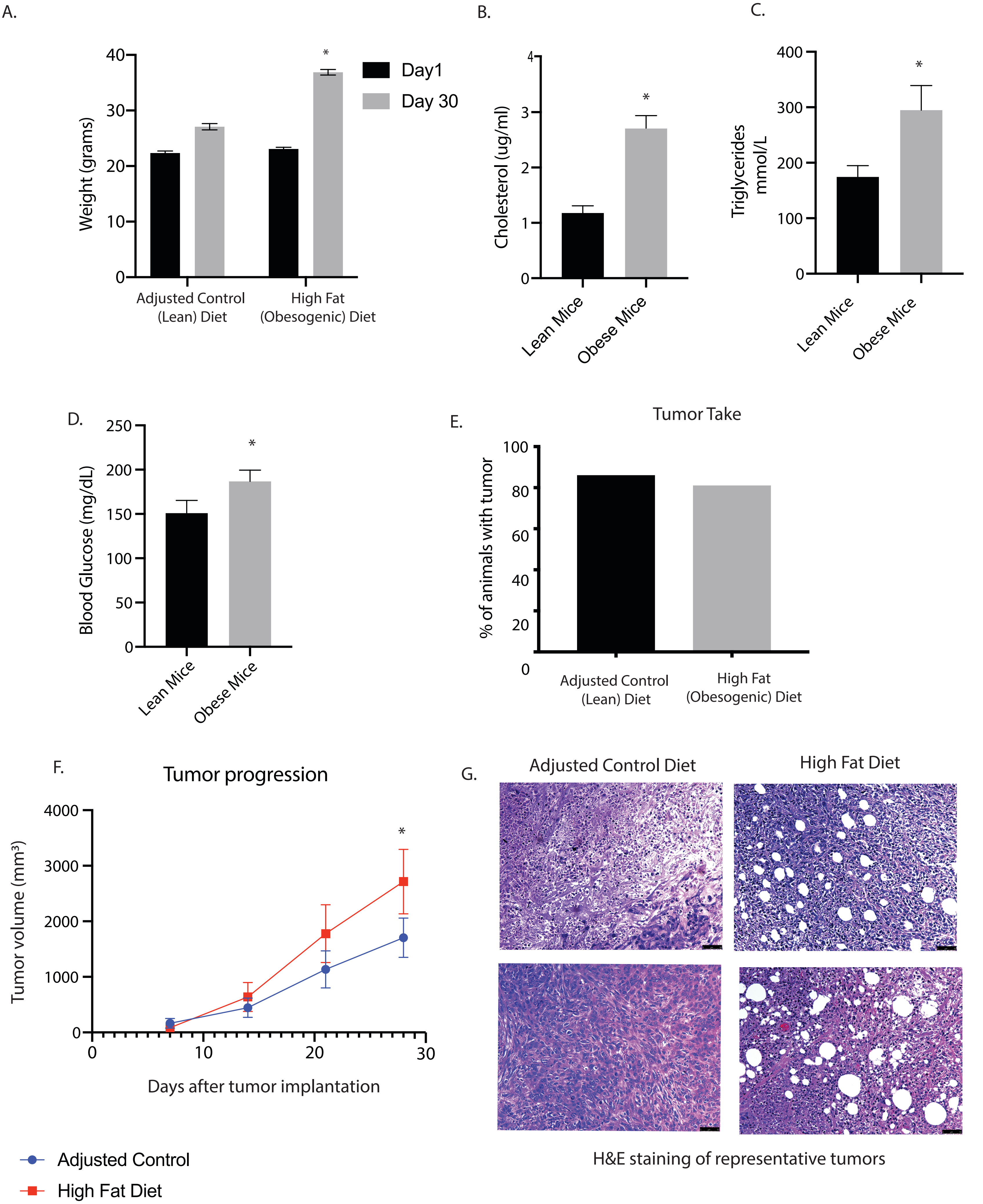

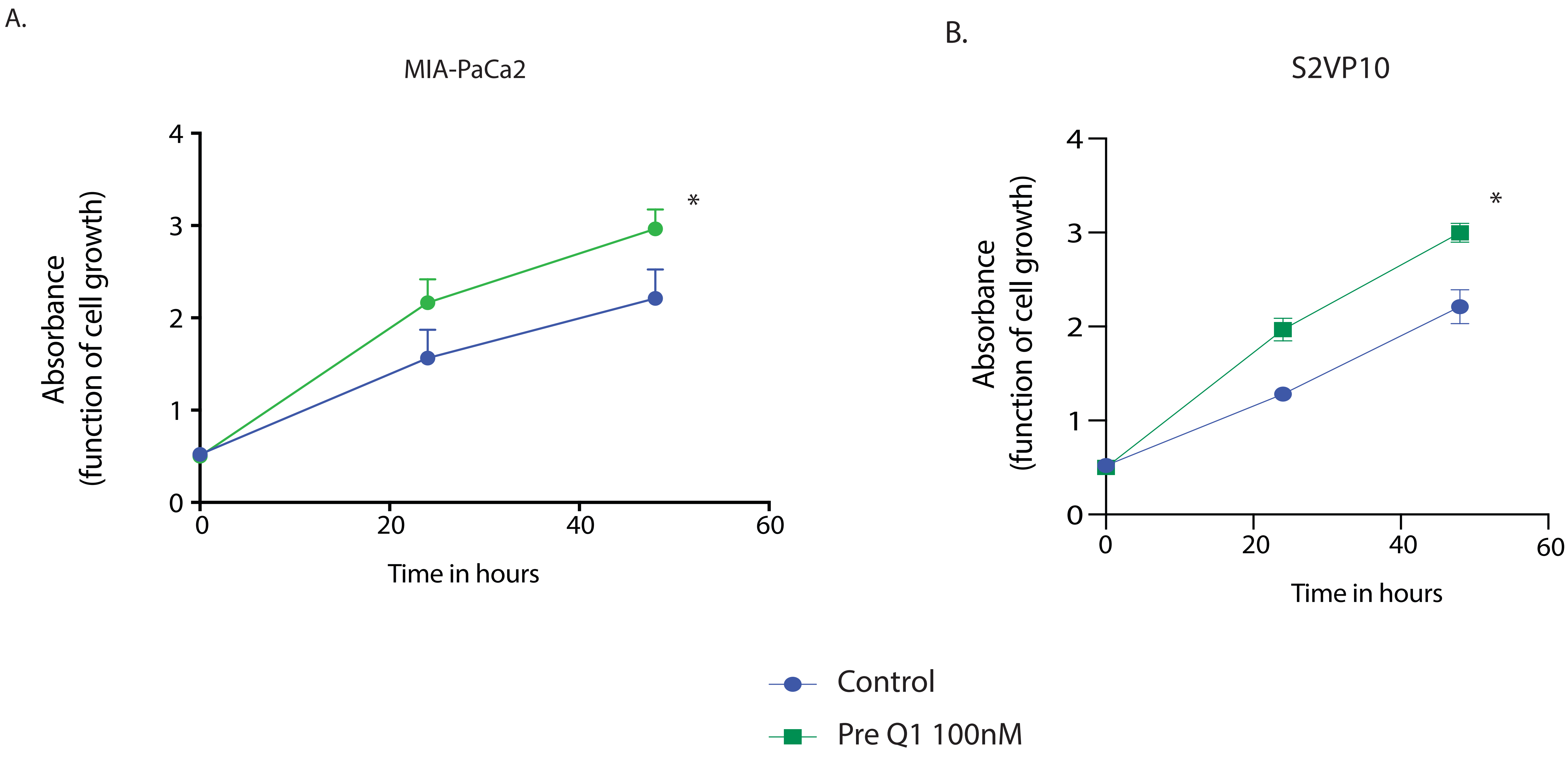

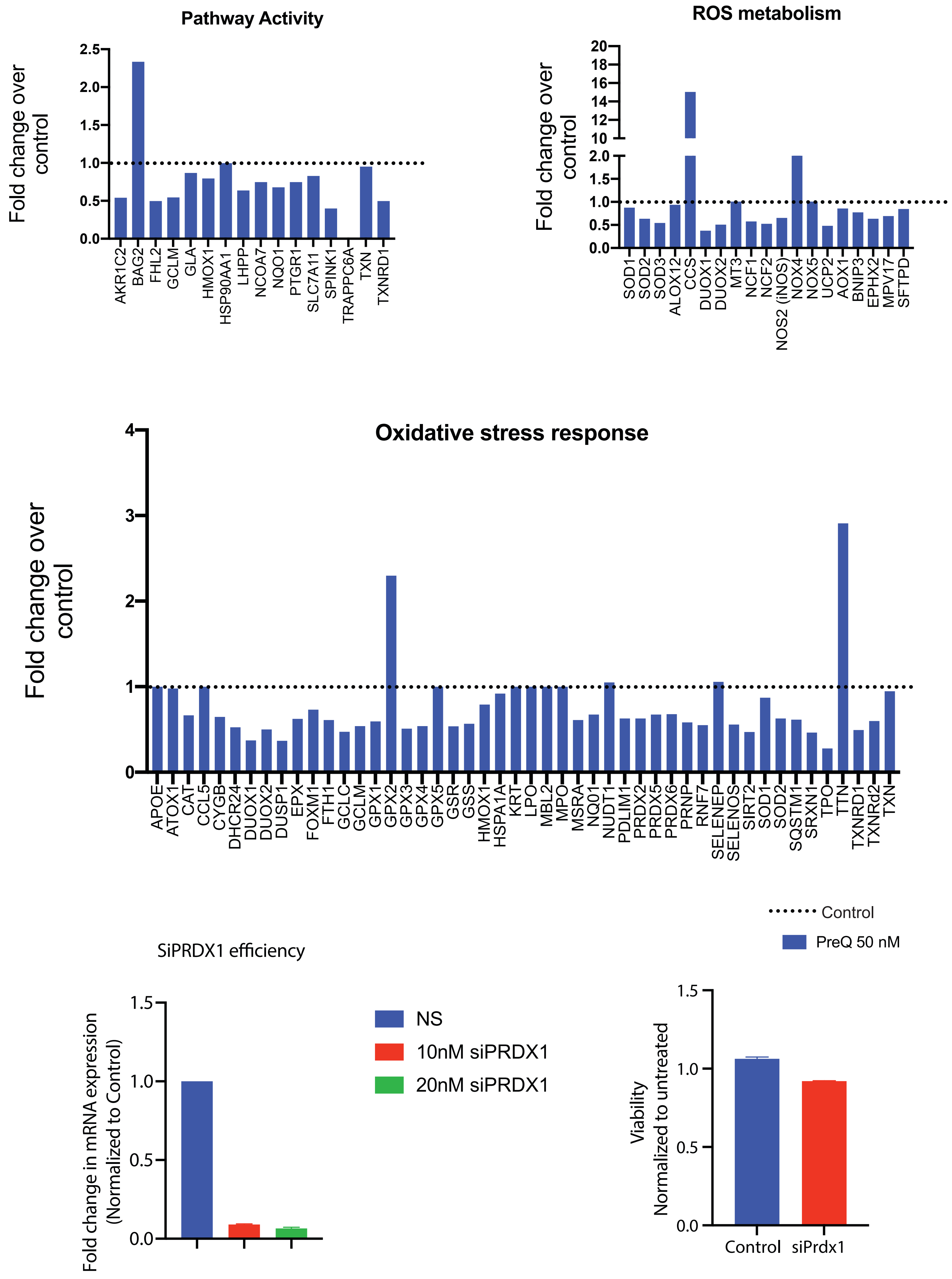

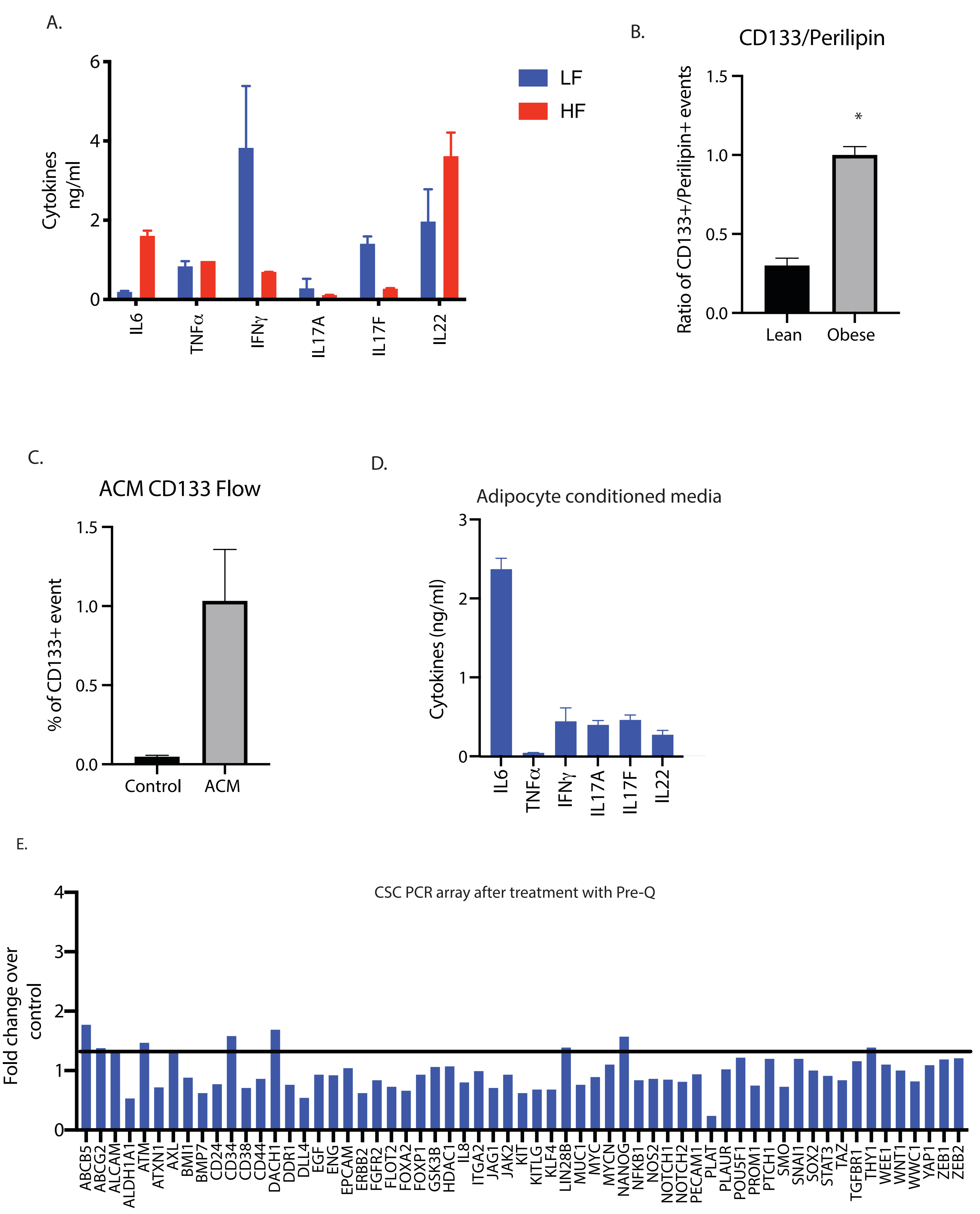
